# Mechanical control of neural plate folding by apical domain alteration

**DOI:** 10.1101/2023.02.10.528047

**Authors:** Miho Matsuda, Jan Rozman, Sassan Ostvar, Karen E. Kasza, Sergei Y. Sokol

## Abstract

Vertebrate neural tube closure is associated with complex changes in cell shape and behavior, however, the relative contribution of these processes to tissue folding is not well understood. In this study, we evaluated morphology of the superficial cell layer in the *Xenopus* neural plate. At the stages corresponding to the onset of tissue folding, we observed the alternation of cells with apically constricting and apically expanding apical domains. The cells had a biased orientation along the anteroposterior (AP) axis. This apical domain heterogeneity required planar cell polarity (PCP) signaling and was especially pronounced at neural plate hinges. Vertex model simulations suggested that spatially dispersed isotropically constricting cells cause the elongation of their non-constricting counterparts along the AP axis. Consistent with this hypothesis, cell-autonomous induction of apical constriction in *Xenopus* ectoderm cells was accompanied by the expansion of adjacent non-constricting cells. Our observations indicate that a subset of isotropically constricting cells can initiate neural plate bending, whereas a ‘tug-of-war’ contest between the force-generating and responding cells reduces its shrinking along the AP axis. This mechanism is an alternative to anisotropic shrinking of cell junctions that are perpendicular to the body axis. We propose that neural folding relies on PCP-dependent transduction of mechanical signals between neuroepithelial cells.

## Introduction

Vertebrate neural tube closure (NTC) is a complex process that involves coordinated behaviors of cells that are controlled by hundreds of genes ^1,2^. Neurulation starts with the specification of neuroectoderm and the formation of the neural plate. The neural plate bends at the medial hinge point to bring about the neural groove and triggers the elevation of the neural folds. The rising neural folds then bend at the paired dorsolateral hinge points and appose each other. Lastly, the folds fuse at the dorsal midline and lose contact with the surface non-neural ectoderm to complete NTC. Failure of NTC is a common birth defect and many responsible genes have been identified, including core components of planar cell polarity (PCP) signaling ^2,3^. Despite extensive research efforts and long-standing debates in the field ^3–6^, elucidation of cellular and molecular mechanisms underlying NTC has been challenging.

Collective cell movements during neurulation are instructed by changes in cell shape and relative positions. Multiple cell behaviors, such as neighbor exchanges, apical constriction and basolateral protrusive activity, have been proposed as the cellular basis for NTC ^7–12^. Whereas early studies suggested that apical constriction is a primary mechanism of tissue folding ^3,13,14^, other reports advocated the importance of neighbor cell exchanges for neural tube elongation ^8,15^. To explain why the neural plate does not shorten in response to isotropic contractions of the apical surface ^4^, cell junctions were proposed to shrink anisotropically in response to PCP signaling ^10^. Alternatively, the elongated shape of the neural tube may be instructed by tissue geometry, location of the constricting cell population or affected by the underlying notochord. Due to divergence of neurulation mechanisms in different species and the complexity of cell shape changes in distinct areas of the neural plate, the relative contribution of various cell behaviors to tissue folding remains poorly understood.

In this study, we evaluated changes in the apical domain of superficial cells in the *Xenopus* neural plate. The neural plate starts bending along specific lines of apically constricted cells at hinge points that are clearly visible in chick embryos ^1,2^ but less pronounced in *Xenopus* ^16^. At early neurula stages, we observed apically constricted cells interspersed with visibly expanded cells that were aligned with the AP axis, especially at the hinge areas. This apical domain heterogeneity required PCP signaling. A mechanical model simulation of the epithelial dynamics showed that the oriented cell elongation of the non-constricting cells can arise as a passive consequence of the presence of constricting cells. *In vivo* analysis of apical domain size in the ectoderm, in which actomyosin contractility has been activated, confirmed that apical constriction and expansion are mechanically coupled. Our analysis proposes that neural plate folding is mediated by PCP-dependent mechanotransduction between neuroepithelial cells.

## RESULTS

### Heterogeneity of apical domains is a hallmark of neural plate folding

Previous time-lapse analyses of *Xenopus* neurulation showed only infrequent cell intercalations in the superficial layer ^8^. This suggests that changes in the neuroepithelial cell shape contribute to neural plate folding more than previously appreciated. *En face* view of phalloidin-stained neurulae showed that the whole neural plate is enriched with F-actin (Fig. 1A), possibly reflecting elevated contractility as compared to the non-neural ectoderm. Binary segmentation revealed variable morphology of apical domains in the cells throughout the neural plate (Fig. 1A, A’). The most pronounced variability concerned the hinge areas of stage 15 neurulae (Fig. 1B, D). Of note, the majority of cells in the neural plate were oriented along the body AP axis (Fig. 1C). We observed that both medial and dorsolateral hinges contained alternating cells with highly constricted apical area as well as the cells elongated along the AP axis (Fig. 1B, D). Quantitative analyses showed increased variability of both apical domain size and cell aspect ratio in the hinge area of the neural plate, compared to those in the non-hinge area and the stage 11 embryonic ectoderm (Fig. 1E, F, Table S1). We conclude that the neural plate acquires considerable cell heterogeneity by the onset of folding, a result supported by another study ^7^. Together, these findings suggest that the observed apical domain heterogeneity plays a role in neural plate folding.

**Figure 1.**
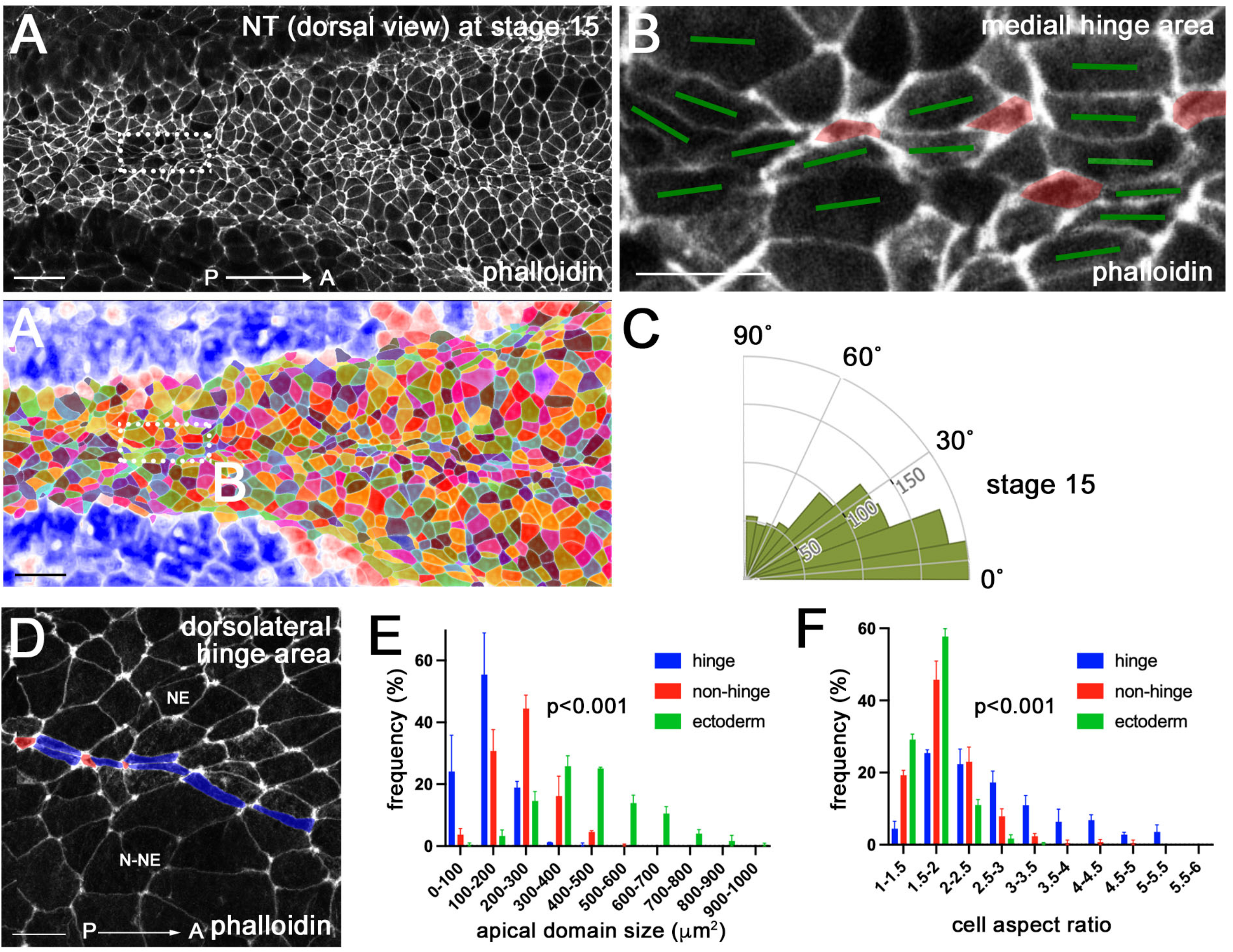
Neural plate hinge cells contain heterogeneous apical domains. (A-B’) Dorsal view of the middle area of a control *Xenopus* neural plate stained with phalloidin (A) and the segmented image (A’) at stage 15. The rectangular area in A, corresponding to the medial hinge, is enlarged in B. (B) Cells with small apical domain (red) are interspersed with large cells elongated along the AP axis (green bars). (C) Rose plots show the orientation of the cells relative to the anteroposterior (AP) axis. n=929. Data from three stage 15 embryos are combined. (D) In the dorsolateral hinge region, cells with small apical domain (red) are adjacent to elongated cells (blue). Neuroepithelium (NE) and non-neural ectoderm (N-NE) are in the upper and lower parts of images, respectively. (E, F) The histogram of apical domain size (E) and cell aspect ratios (E) of cells in stage 11 embryonic ectoderm (green), and cells in hinge (blue) and non-hinge (red) areas of stage 15 neural plate, Data (means +/- s.d.) are combined for three embryos, representative of three independent experiments. NT hinge, n=255; NT non-hinge n=426; stage 11 ectoderm, n=1218. Coefficients of variation (CV) are shown in Table S1. Statistical significance was assessed using Kolmogorov-Smirnov test. Scale bars are 50 μm in A, A’, 20 μm in B and D.

### PCP signaling is required for apical domain heterogeneity in the neural plate

Coordinated cell polarization in the epithelial plane, known as planar cell polarity (PCP), commonly underlies cell shape changes during morphogenesis, including vertebrate NTC ^17–20^. PCP in the neural plate can be visualized by nonuniform distribution of core PCP protein complexes and requires the function of the conserved core PCP protein Vangl2 ^21,22^. Our previous studies suggested that PCP signaling is required for apical constriction in the blastopore during *Xenopus* gastrulation and during NTC ^23,24^. Therefore, we asked whether PCP signaling regulates the apical domain heterogeneity in the neural plate. Unilateral knockdown of *vangl2* using previously characterized *vangl2* morpholino oligonucleotide (MO)^23,25^ with GFP as a lineage tracer inhibited apical domain heterogeneity in the MO-injected half of the neural plate (Figs. 2A-B’, S1A-B’). In addition, F-actin enrichment in the control side of the neural plate was reduced at the MO-injected side (Fig. 2B, B’). Of note, the effect of *vangl2* knockdown was visible in both posterior (Fig. 2B-B’) and anterior neural plate (Fig. S1C-E), but absent in the non-neural ectoderm (epidermis, N-NE)(Fig. S1F, G). Similar observations were made for the knockdown of *fuzzy* (data not shown), another conserved protein required for PCP in *Drosophila* ^26^ and NTC in *Xenopus* ^27^. Our findings indicate that the cell heterogeneity develops in the neural plate under the control of PCP signaling.

**Figure 2.**
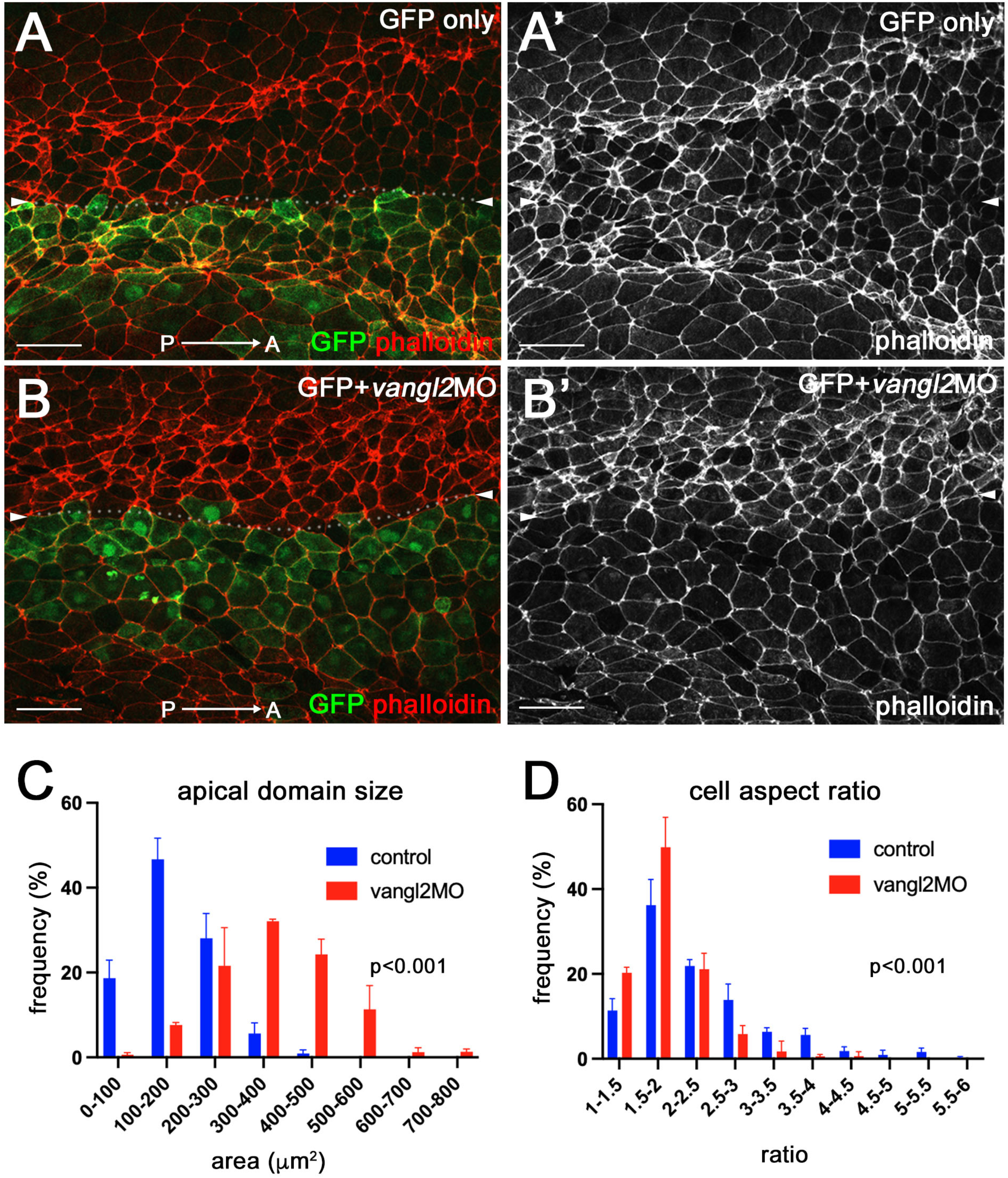
PCP signaling is required for apical domain heterogeneity in the neural plate. (A-B’) Representative images of control GFP-injected embryo and *vangl2* MO-injected embryo (the injected side marked by GFP). Dorsal view. Control GFP RNA (A, A’), or GFP RNA and 10 ng *vangl2* MO (B, B’) were injected into four-cell embryos targeting the presumptive neural plate. Control, n=6; *vangl2* MO, n=9. (C, D) The histogram of apical domain size (C) and cell aspect ratio (D) of uninjected and *vangl2* MO-injected cells in B, B’. Data (means +/- s.d.) was quantified for three embryos in each group and represent three independent experiments. Statistical significance was assessed using Kolmogorov-Smirnov test. CVs are shown in Table S1. Scale bars are 50 μm in A-B’.

### Cell elongation can arise passively from presence of interspersed contractile cells

The heterogeneity of apical domains in the neural plate indicates that only a subset of the cells undergo apical constriction. We asked whether the observed cell behavior in the tissue can be explained as a mechanical consequence of apical contractility. We therefore analyzed a 2D vertex model ^28–30^ of the neural plate epithelium. In a 2D vertex model, each cell is represented as a polygon and defined by the position of its vertices. An energy is chosen for the model tissue, and the vertices are then assumed to follow overdamped dynamics towards an energy minimum. In this case, we use the usual area-and-perimeter elasticity energy, which sets a target area and perimeter value for each cell ^28^. The energy therefore reads

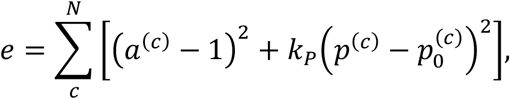

where the sum runs over all *N* cells, *a*^(*c*)^ and *p*^(*c*)^ are the area and the perimeter, respectively, of the *c*-th cell, whereas 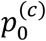 is its target perimeter; *k_p_* is the perimeter elasticity modulus (this is the dimensionless form of the energy; see Methods). We set 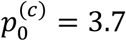, just below the perimeter of a regular hexagon of area 1, and started the simulation with a regular hexagonal lattice. A *N_x_* by *N_y_* rectangular region (blue border in Fig. 3A) represents the posterior region of the neural plate. Here, *N_x_* corresponds to the length along the AP axis, and *N_y_* the length perpendicular to the AP axis. Unless stated otherwise, we set *N_x_* = 60 and *N_y_* = 20.

**Figure 3.**
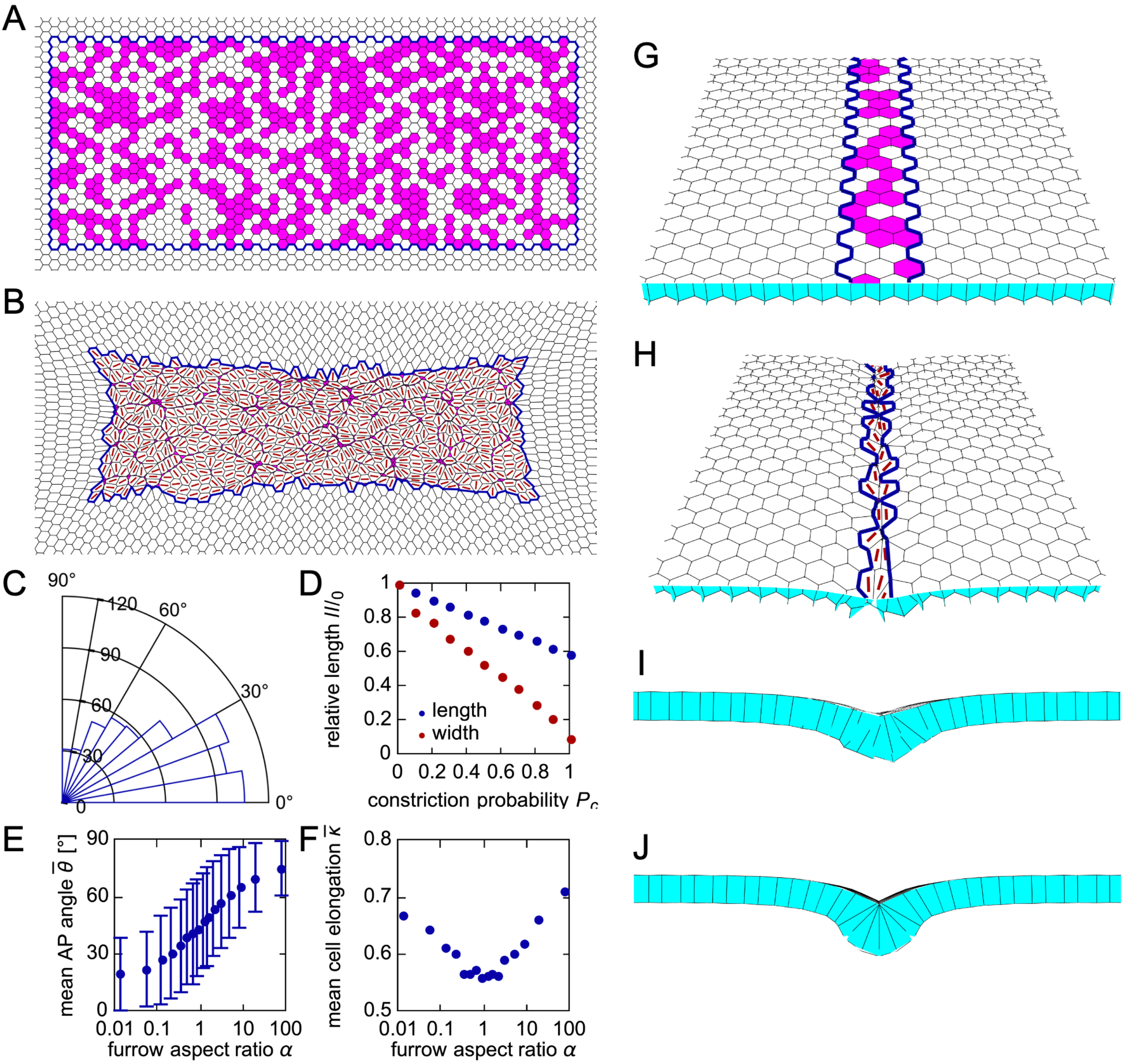
Mechanical model shows that contractile cell subpopulation leads to cell elongation and alignment. (A-F) 2D vertex model of the neural plate. (A) Zoom on the neural plate region of the vertex model initial condition. Blue border outlines the posterior neural plate region of the model tissue. Apically constricting cells are in magenta (tissue shown for *P_c_* = 0.5). (B) Model tissue from panel A after relaxation at *t* = 2000. Red lines show the direction of cell elongation. (C) Distribution of angles between the AP axis (horizontal line) and cell elongation direction of non-constricting neural plate cells for the model tissue in panel B; 0° corresponds to perfect alignment. (D) Central length (along AP) and width (perpendicular to AP) of the model tissue after relaxation as a function of the probability of cell constriction *P_c_*; values are normalized by length and width if no cells constrict. (E) Mean angle with the AP axis as a function of the constricting region aspect ratio *α*. Error bars indicate the standard deviation between non-constricting cells in a model tissue. (F) Mean cell elongation as a function of the furrow aspect ratio. See Methods for details. (G-J) Furrow formation in a 3D vertex model simulation of the hinge area. (G) Initial condition for the 3D vertex model simulation. Hinge region is outlined by a blue border and constricting cells are shown in magenta (tissue shown for *P_c_* = 0.5). (H) Model tissue from panel G after relaxation at time *t* = 5000. Red lines show the direction of elongation for the apical side of the cells. (I) Cross-section view of the model tissue in panel H, showing the formation of a shallow furrow. (J) Cross-section view of a model tissue with *P_c_* = 1.

At the start of the simulation, each cell in the neural plate region has a probability *P_c_* of becoming apically constricting (Fig. 3A). We modeled constriction by setting the target perimeter of constricting cells to 10% of that of the non-constricting cells. Fig. 3B shows the tissue shape after relaxation at time *t* = 2000, with red lines indicating the direction of elongation for the non-constricting cells, whereas Fig. 3C shows the distribution of angles between cell elongation and the AP axis. This simulation suggests that the anisometric shape of the constricting region is sufficient for the non-constricting cells to favor elongation along the region’s long axis. If only a fraction of cells in the neural plate constrict, the tissue shrinks less than when all cells constrict (Fig. 3D). Note that in the model, the non-constricting cells elongate but do not usually expand; their final average area is ~9% lower than in the initial condition.

The underlying physical mechanism for cell elongation is therefore similar to the one proposed to explain why the mesodermal domain during *Drosophila* ventral furrow formation contracts much less along the AP axis than perpendicular to it ^31^. A rectangular contractile domain in a flat 2D elastic plate will contract more along the short than the long axis of the domain ^32,33^. A similar mechanism has recently also been studied in 3D ^34^. Importantly, this also implies that the external tissue outside the neural plate counterbalancing the deformation is an important factor in the observed cell elongation ^32^. The difference in the present model when compared to the *Drosophila* models above is that there are now two populations, with only one generating the contractility. A related behavior has also been reported and modeled during the slow phase of *Drosophila* ventral furrow formation ^35,36^.

We also ran simulations with different *N_x_* and *N_y_*, changing the aspect ratio of the constricting region *α* = *N_y_/N_x_*, but keeping the number of cells in it similar to the above case. We found that cells are on average aligned closer to the *x* axis for *α* < 1, whereas they align closer to the *y* axis for *α* > 1 (Fig. 3E). As the anisometry of the region (i.e., aspect ratio either increasing or falling away from 1) increased, cells also became more elongated (Fig. 3F).

We next considered a more detailed model that includes a higher fraction of constricting cells in the two hinges. We separated the neural plate into two 3-cell-wide regions, representing the hinges, which are separated by ~14 cells. Cells in the hinges have a probability *P_h_* = 0.5 of constricting, whereas those in the region between hinges only constrict with a probability *P_c_* = 0.2 (Fig. S2A). Interestingly, we found that in this case only cells in the hinges elongate along the AP axis, whereas the remaining cells in the neural plate do not have a notable preferred direction of elongation (Fig. S2B-D). Thus, the geometry of the shrinking tissue depends on the frequency and distribution of apically constricting cells.

Lastly, we used a 3D vertex model ^37–39^(Fig. 3G-J, see Methods) to test whether the constriction of only a fraction of cells is still sufficient for the formation of the furrow. Here, we are only interested in the hinge, so we limited the region in which cells have a probability of constricting to a ~3-cell-wide patch (blue border in Fig. 3G), with periodic boundary conditions along the AP axis. As in the 2D model, constriction was modeled by reducing the target perimeter of the apical cell side to 10% of its initial value. The resulting tissue morphology after relaxation at *P_c_* = 0.5 is shown in Fig. 3H and I: the constriction of only ~50% of cells is indeed sufficient for the formation of at least a shallow furrow. However, the furrow is deeper in a model tissue in which all hinge cells constrict (Fig. 3J). Altogether, our simulations allow us to propose that cell elongation in the neural plate is a passive mechanical consequence of cell constriction. Lastly, we chose to focus on minimal models of the interplay between cell constriction and the consequent elongation. Therefore, additional mechanisms included in other vertex models of neural tube formation ^40,41^ are not considered here.

### Lmo7-expressing ectoderm as a model of apical domain heterogeneity

Our *in silico* model predicts that mechanical forces are sufficient to instruct cell elongation and orientation in a tissue with a subset of dispersed contractile cells. Testing this hypothesis in the neural plate is challenging due to its complex tissue dynamics and patterning events. We therefore examined apical domain changes in constricting gastrula ectoderm, a tissue from which the neural plate originates.

We induced apical constriction in *Xenopus* ectoderm using Lmo7, a myosin II-interacting protein, as we recently reported ^42^. The injection of *gfp-lmo7* RNA in two opposing blastomeres of 4-8-cell embryos resulted in ectoderm hyperpigmentation (Fig. 4A, B) that commonly accompanies apical constriction ^42^. Notably, the majority (80-90%) of injected embryos showed coordinated cell movements (Movie 1) and developed a pigmented furrow between the two injection sites (Fig. 4B, C). Importantly, GFP-Lmo7 cells in the furrow were aligned (Fig. 4D, E). Similar hyperpigmentation and furrow formation have been observed in the ectoderm cells after the induction of apical constriction by the actin-binding protein Shroom3 ^43,44^. Since both Lmo7 and Shroom3 induce apical constriction and participate in NTC, the cellular mechanisms underlying furrow formation in the apically constricting ectoderm may be akin to those operating in the neural plate. We therefore used apical constriction in ectoderm to study the processes underlying neural plate folding.

**Figure 4.**
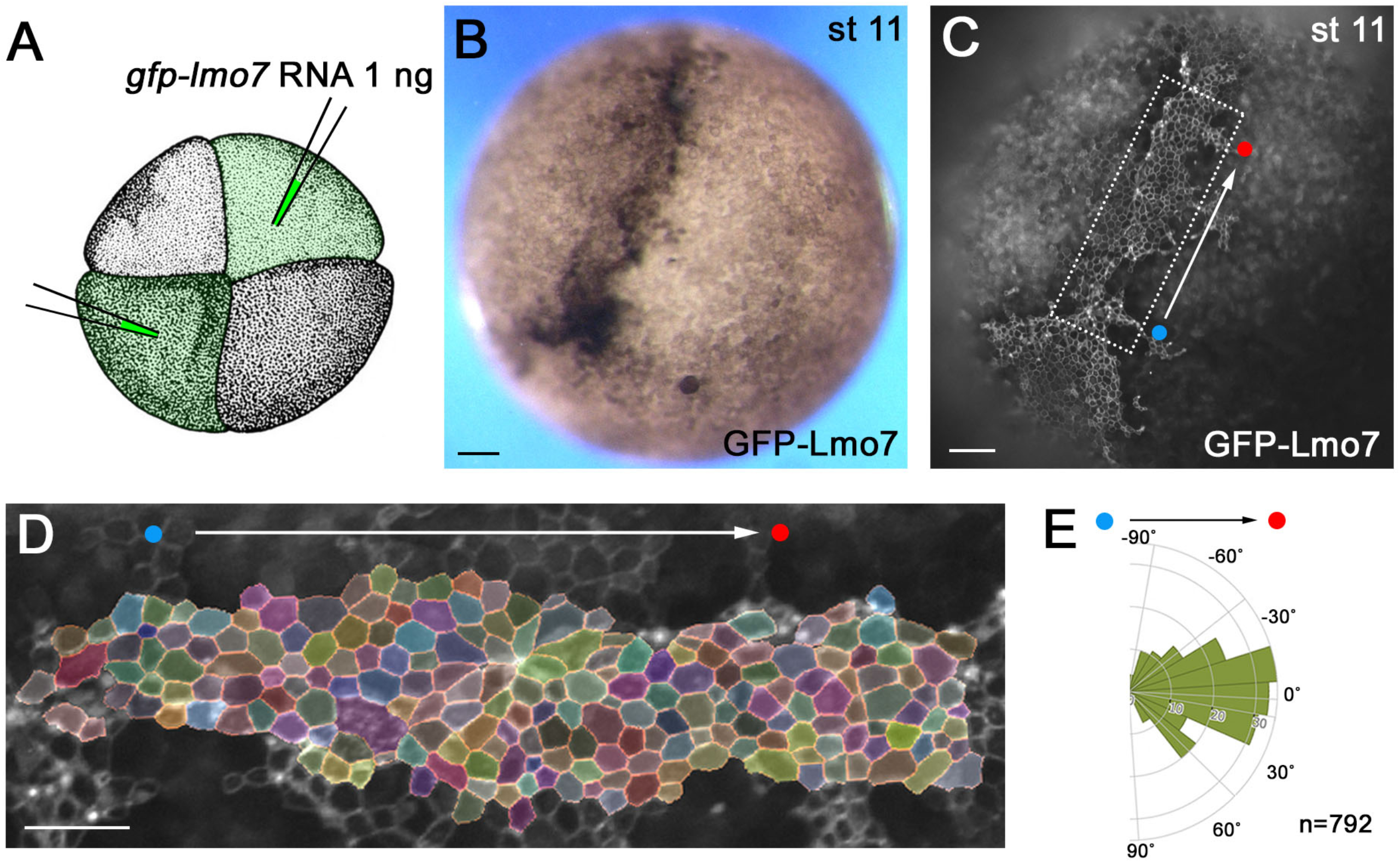
Lmo7-expressing ectoderm as a model of apical domain heterogeneity. (A) Scheme of the experiment. GFP-Lmo7 RNA (1 ng) was injected in two oppositely localized animal blastomeres in 4-cell stage embryos. (B) Representative GFP-Lmo7-expressing embryo at stage 11. Hyperpigmentation was observed at 90-95% penetrance (n>100). (C) Still image from Movie 1 shows GFP-Lmo7 fluorescence from the embryo in B. (D) Segmented image of the rectangular area in C. (E) Representative rose plot from 792 cells from one embryo depicts cell orientation with respect to the injection axis (white arrow, red and blue dots in C-E). The data represent at least three experiments. Scale bars are 100 μm in B, C and 50 μm in D.

### Apical domain dynamics in Lmo7-expressing ectoderm

We monitored apical domain dynamics by live imaging of embryos microinjected into the four animal blastomeres with *gfp-lmo7* RNA (100-200 pg) (Fig. 5A). At this dose, GFP-Lmo7 was uniformly expressed in the ectoderm and the expressing cells had similar apical domain size at the beginning of imaging at stages 10.5-11 (Fig. 5B). By the end of imaging (3.5-4 hrs), however, some cells apically constricted (AC cells, red cells in Fig. 5B-D’) whereas others exhibited continuous apical expansion (AE cells) (blue cells in Fig. 5B-D’)(Movie 2). We confirmed this observation by quantifying apical domain size dynamics of individual cells in several GFP-Lmo7 embryos (Figs. 5E, G, S3A-B””, S4B, C). Of note, AC cells were often located adjacent to AE cells (Fig. 5B-D’). By contrast, apical domain size did not significantly change in control cells that expressed GFP-ZO1 (Figs. 5F, G, S3C-D”’)(Movie 3) or membrane-tethered GFP-CAAX (Fig. S4A, C).

**Figure 5.**
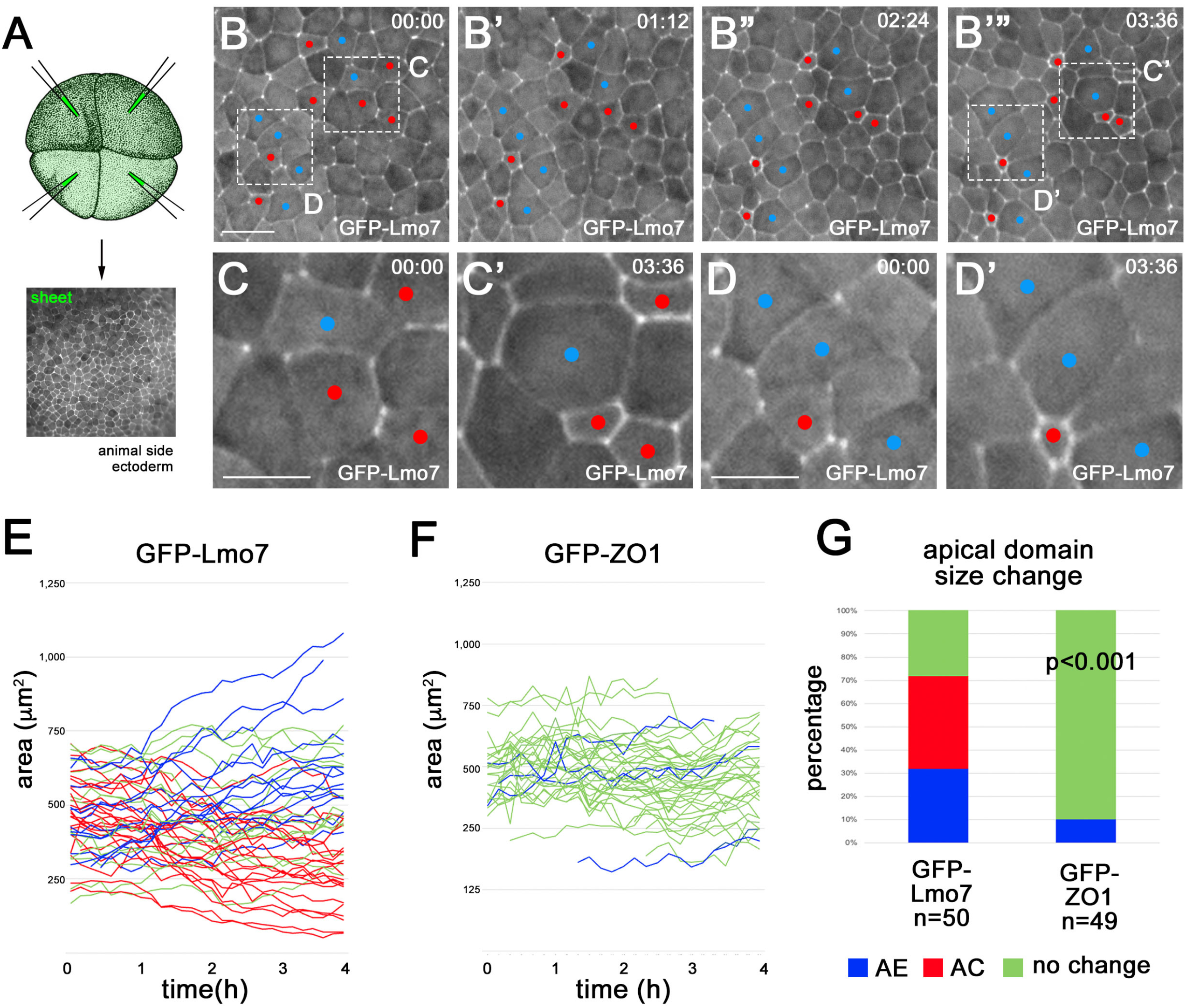
Apical domain heterogeneity in ectoderm cells expressing Lmo7. (A) Scheme of the experiment. GFP-Lmo7 or GFP-ZO1 RNA (150 pg) was injected into four animal blastomeres of 4-8-cell embryos for live imaging at stage 11. Uniform fluorescence has been confirmed in stage 11 ectoderm. (B-C”) Time-lapse imaging of GFP-Lmo7 embryos for approximately 4 hrs. Apically constricting (AC) and expanding (AE) cells are marked by red and blue, respectively. Areas in B-B” are enlarged in C-D”. (E, F) Quantification of apical domain dynamics in GFP-Lmo7 (E) and GFP-ZO1 (F) cells in one representative embryo. Each line represents apical domain size changes of individual cells over 4 hrs. The cells were scored as AC (red) or AE (blue) if they had more than 20% decrease or increase in their apical domain size, respectively. The remaining cells were designated as ‘no change’ (green). GFP-Lmo7 (n=50). GFP-ZO1 (n=49). (G) Frequencies of cells with different changes in apical domain size are shown for samples in E and F. p<0.001. Statistical significance was assessed using the Freeman-Halton extension of Fisher’s exact test.. These experiments have been repeated 3-5 times. Scale bars are 40 μm in B and 20 μm in C, D.

These observations demonstrate the emergence of the striking heterogeneity of apical domains in Lmo7-expressing epithelium that is similar to the one visible in the neural plate.

### Cells mosaically-expressing Lmo7 are predominantly constricting

Uniform Lmo7 expression in *Xenopus* ectoderm triggered the appearance of both AC and AE cells. This observation may be explained by direct effects of Lmo7 on apical constriction and apical expansion. Alternatively, AE cells may form as a passive response to the AC cells as predicted by our vertex model. To test these possibilities, we compared apical domain dynamics in embryos expressing Lmo7 uniformly (‘sheet’ in Fig. 5A) or mosaically (‘isolated’, Fig. 6A, S5A). We confirmed that the uniform expression of Lmo7 triggered both apical constriction and expansion (Fig. 6B-C’, H, J)(Movie 4), whereas the majority of ‘mosaic’ GFP-Lmo7 cells were constricting (Fig. 6D-G’, I, J, S5C-C”, F, G)(Movie 5). Note that due to a shorter time-lapse imaging period, the frequencies of AC and AE cells are smaller as compared with 4-5 hr imaging (compare Figs. 5G, S4C and Figs. 6J). By contrast, control cells expressing myr-BFP did not show a significant change in the apical domain size (Fig. S5D-D”, E, G). These results suggest that the primary function of Lmo7 is to induce apical constriction. Since the AE cells were often found adjacent to the AC cells (Figs. 5B-6D’”, 6B-C’), the increase of their apical surface was likely driven by the pulling force of neighboring AC cells.

**Figure 6.**
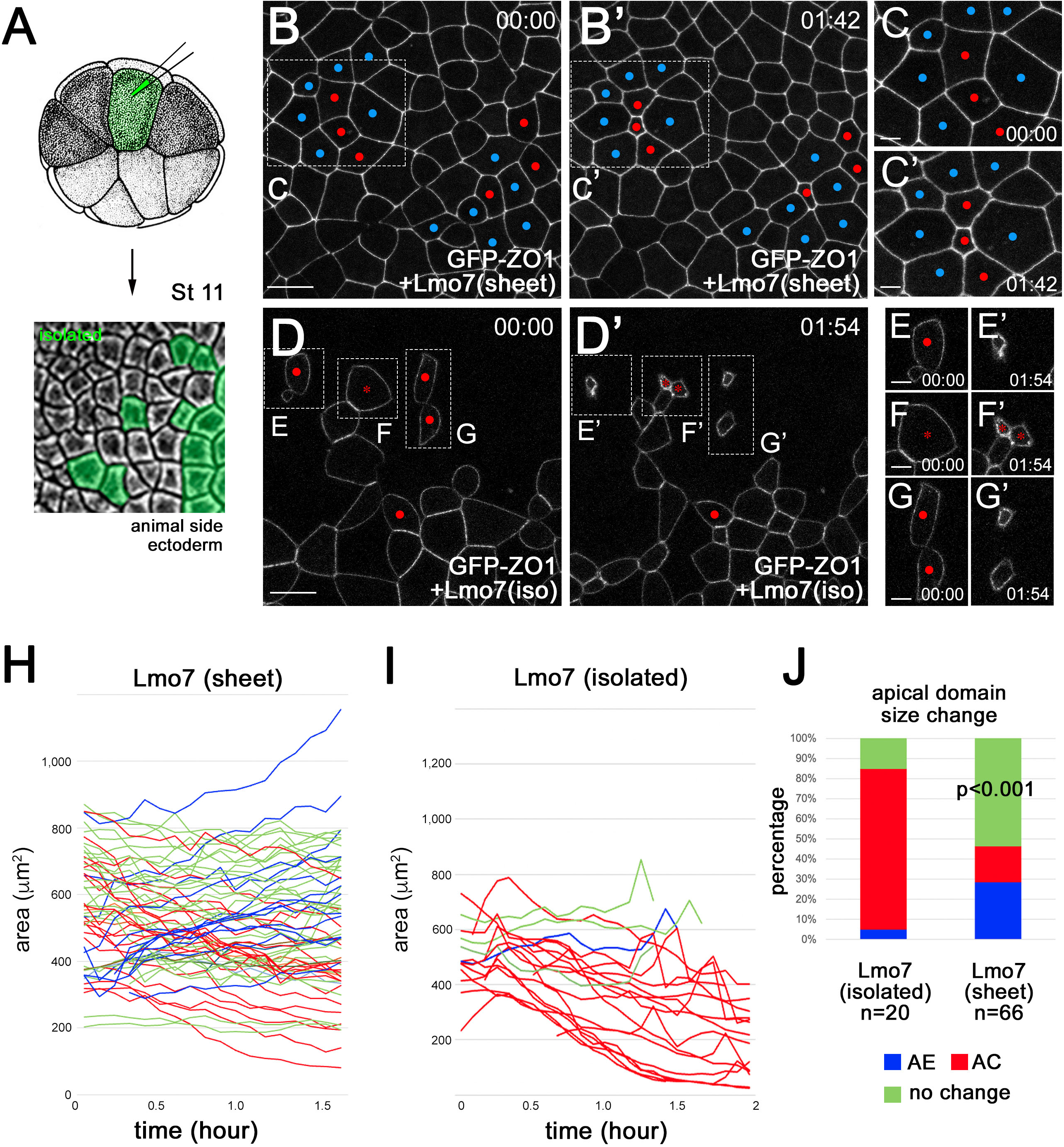
Mosaic expression of Lmo7 causes apical constriction. (A) Scheme of the experiment. GFP-ZO1 RNA (100 ng) and Flag-Lmo7 RNA (150 pg) were coinjected into one ventral blastomere of 16-cell stage embryos. (B-G’) Still images of the “sheet” (B-C’, Movie 2) and “isolated” (D-G’, Movie 3) cells in GFP-Lmo7 embryos. Time-lapse imaging was initiated at stage 11. (D-G’). AC and AE cells are marked by red and blue, respectively. Rectangular areas in B,B’ and D,D’ are enlarged in C,C’ and E-G’, respectively. Scale bars are 30 μm in B, D and 10 μm in C, E-G. (H, I) Apical domain dynamics in the “sheet” (H) and “isolated” ectoderm (I). Each line represents apical domain size changes of an individual cell over 1.5-2 hours. AC, AE and ‘no change’ cells (See Fig. 5 legend) are shown in red, blue and green, respectively. Quantification was based on n=66 cells from one GFP-Lmo7 (sheet) embryo and n=20 cells from three GFP-Lmo7 (isolated) embryos. (J) Frequencies (%) of AC, AE and ‘no change’ cells in H and I. These experiments have been repeated 3-5 times. p<0.001, the Freeman-Halton extension of Fisher’s exact test.

### Apical domain heterogeneity in ectoderm cells expressing Shroom3

We next asked whether the observed modulation of apical domain size was specific to Lmo7 or shared by other apical constriction regulators. Shroom3 is a known to induce apical constriction and is essential for vertebrate NTC ^43,44^. Similar to Lmo7 cells, the ectoderm cell population uniformly expressing Shroom3 had increased aspect ratio (Fig. S6) and were oriented relative to the axis connecting two injection sites (line of constriction, Fig. S6C).

We also examined apical domain size in ectoderm cells mosaically-expressing Shroom3 (Fig. 7A). We observed that these cells had smaller apical domains (Fig. 7B-D) and a wider range of cell aspect ratio compared to adjacent wild-type cells (Fig. 7E). These cells (blue, Fig. 7C, C’) adjacent to Shroom3 cells (red, Fig. 7C, C’) were oriented relative to the line of constriction (green lines in Fig. 7C’). These findings support the view that the coordinate regulation of apical domain size is a general property of epithelial morphogenesis.

**Figure 7.**
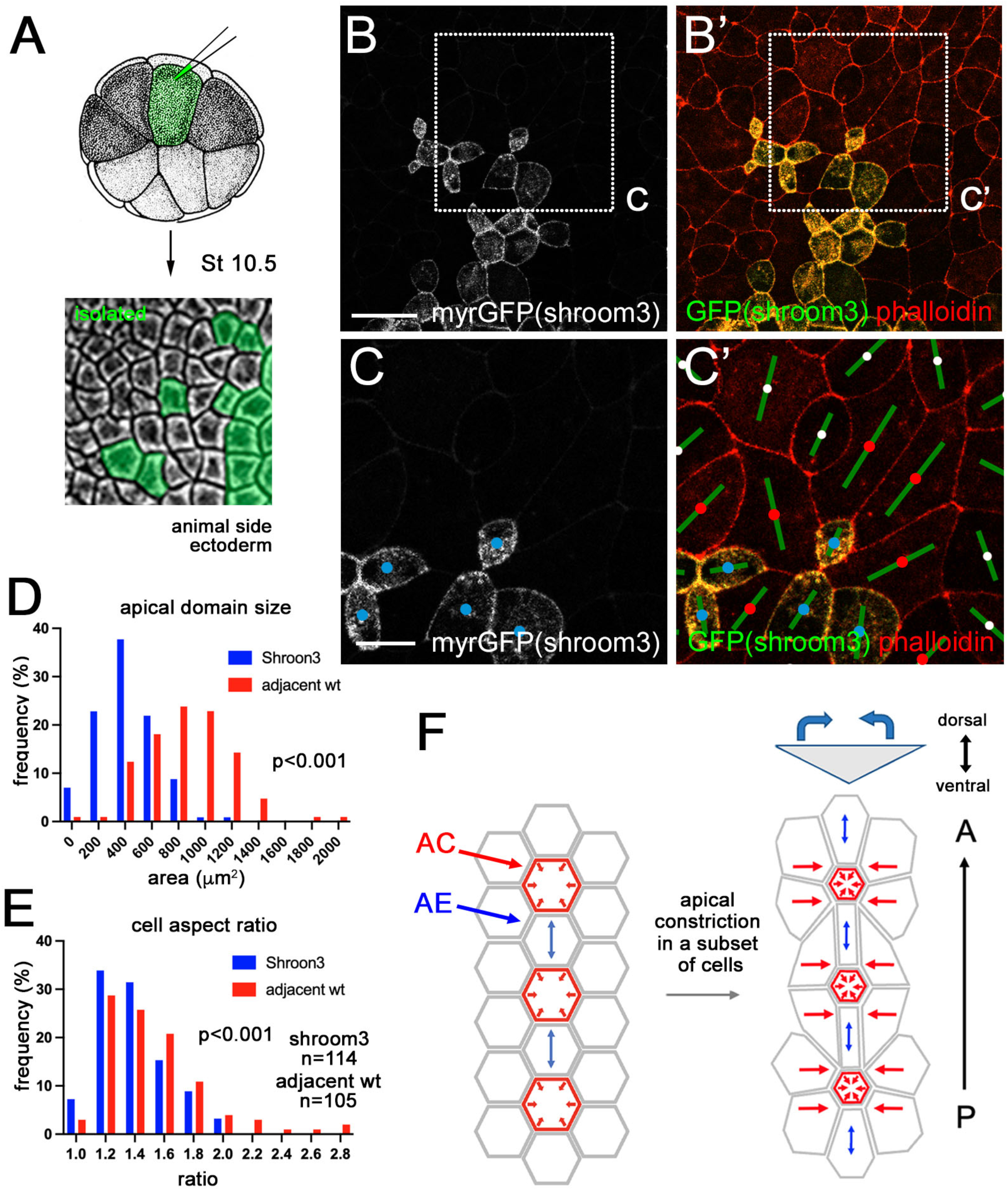
Modulation of the apical domain by Shroom3. (A) Scheme of the experiment for G-I. Shroom3 (80 pg) and myrGFP (50 pg) RNA were co-injected into one animal blastomere of 16-cell stage embryos. Stage 10.5 embryos were fixed, stained with phalloidin and the animal pole ectoderm was imaged. (B-C’) Shroom3 cells (red) with adjacent wildtype cells (blue) are shown. Rectangular areas in B and B’ are enlarged in C and C’. Cell orientation is shown by green bars. (D) Apical domain size of Shroom3 cells and the neighboring wild-type cells. (E) Cell aspect ratios for Shroom3 cells and the adjacent cells. n=114 (Shroom3 cells) and n=105 (adjacent wild-type cells) from 5 embryos. p<0.001, the Kolmogorov-Smirnov test. (F) Model. Tissue bending is driven by isotropic apical constriction of a subset of cells (red arrows) at the hinge areas of the neural plate. Due to the geometry of the neural plate, neighboring cells passively respond by elongating along the anteroposterior axis (blue arrows). This behavior promotes neural plate folding and preserves tissue length. Scale bars are 50 μm in B and 20 μm in C.

Based on our observations, we propose a model in which the dispersed population of isotropically constricting cells is sufficient to drive neural plate bending, whereas the coexistence of the shrinking and elongating cells at the hinges reduces tissue shrinking along the AP axis (Fig. 7F).

## Discussion

This study investigated the origin of apical domain heterogeneity in the vertebrate neural plate and its contribution to NTC. Although mediolateral and radial cell intercalations can contribute to neural tube elongation, this mechanism has not been supported by our direct observations as only rare cell intercalations have been observed in the neural plate at stages 13-15. By contrast, apical constriction is manifested throughout the length of the neural plate, most notably at the medial and dorsolateral hinges ^7,16,45^, consistent with its being a driver of neural plate bending. We observed that apically constricting cells were interspersed with cells that apically expanded along the AP neural axis, especially in hinge areas. Simulations of the tissue constriction using 2D and 3D vertex models fit well with our experimental observations. To further understand the mechanisms underlying cell shape changes, we monitored apical constriction induced in the superficial ectoderm by Lmo7 and Shroom3. Experimental induction of AC and AE cells in the ectoderm caused cell alignment relative to the apical constriction axis and triggered furrow formation, which suggests that apical domain heterogeneity contributes to epithelial folding. We find that a subpopulation of dispersed contractile cells is sufficient to trigger epithelial bending and that the apically expanded cells react to the pulling forces of their constricted neighbors. Thus, the elongated shape of cells at the hinges may be driven by passive mechanical signals instead of planar polarized junctional shrinking. Our observations highlight similar roles of mechanical forces in the regulation of cell shape and epithelial tissue dynamics in other systems, especially in *Drosophila* ventral furrow formation ^31,46–49^.

Early studies argued that isotropic apical surface contraction should cause shortening of the neural tube, contrasting the observed elongation of the anteroposterior body axis ^4^. We propose that the passive reaction of non-constricting cells to the tensile forces of their constricting neighbors causes apical domain expansion and alignment along the AP axis (Fig. 7F). Thus, the presence of these two cell morphologies allows the neural plate to fold at the hinges while maintaining its overall length. We suggest this mechanism as an alternative to other models of NTC ^3–6,40^. According to the Takeichi model, NTC in the chick embryo involves PCP-dependent anisotropic shrinking of the cell junctions that are perpendicular to the AP axis ^10^. In our model, the force-generating cells are mechanically isotropic, but the forces they produce expand apical domain size and aspect ratio of the adjacent cells. Although the outcome of both models is the same, we expect that the contractile behavior is activated only in the AC cells, while the AE cells passively follow the morphological changes of their neighbors. Consistent with our model, neural tissue deformations are more significant after laser cutting of the cell junctions that are along the AP axis as compared to the ones perpendicular to the axis ^50^. To distinguish the two models, further studies are needed to compare the subcellular localization of various contractility markers.

The morphogenetic behaviors that we described in the neural plate (and, presumably, the underlying physical forces) require intact PCP signaling because they were inhibited by the knockdown of *vangl2*, a core PCP protein. This finding supports earlier studies connecting PCP proteins to apical constriction ^23,51^. Of note, we observed the effects of PCP proteins on both anterior and posterior regions of the neural plate, at variance with other recent reports ^7,45^. This difference might be explained by varying degree of PCP protein depletion. Given that the expanding cells respond to neighboring cell contractility, we expect that the effect of PCP on these cells is indirect.

Several possibilities may be considered to explain the origin of apical domain heterogeneity in the neural plate. First, global patterning factors, such as neural tissue inducing BMP inhibitors ^52^, should play a role, because apical constriction is not normally observed in non-neural ectoderm and because the position of dorsolateral hinges tightly correlates with the neural plate border. Second, the ‘salt-and-pepper’ pattern of cells with large and small apical domains may form stochastically in response to slight variations of contractile activity. Third, the location of the cells with a particular morphology may be transcriptionally controlled and amplified by lateral inhibition, e.g. by the Notch pathway ^53^. Additional studies are warranted to explain how the contractility of the neural plate is controlled and why actively constricting areas often contain more than a single cell. Whether the morphological changes observed in our study are subject to mechanical, transcriptional or post-transcriptional regulation, they likely play critical roles in many diverse tissues and processes, including *Drosophila* ventral furrow formation ^54^ or *Xenopus* blastopore ^55^.

## Methods

### Xenopus embryos, plasmids and microinjections

Wild-type *Xenopus laevis* were purchased from Nasco and Xenopus1, maintained and handled following the Guide for the Care and Use of Laboratory Animals of the National Institutes of Health. A protocol for animal use was approved by the Institutional Animal Care and Use Committee (IACUC) at Icahn School of Medicine at Mount Sinai. *In vitro* fertilization and embryo culture were performed as described ^24^. Embryo staging was determined according to Nieuwkoop and Faber ^56^. For microinjections, embryos were transferred into 3% Ficoll 400 (Pharmacia) in 0.5x Marc’s modified Ringer’s (MMR) solution (50 mM NaCl, 1 mM KCl, 1 mM CaCl2, 0.5 mM MgCl2 and 2.5 mM HEPES (pH 7.4) ^57^.

Plasmids encoding GFP-Lmo7, Flag-Lmo7 and GFP have been previously described ^42^. pCS2-myr-tagBFP, -GFP or -RFP were generated by PCR of tagBFP, GFP or RFP and annealing of chemically synthesized oligonucleotides encoding a myristoylation site. pCS2-3xGFP-mZO1 contains in-frame fusion of three GFP open reading frames and mouse ZO1 cDNA. Details of cloning are available upon request. Capped RNA was synthesized using mMessage mMachine SP6 Transcription kit (Invitrogen) and purified by LiCl precipitation or RNeasy mini kit (Qiagen). RNA in 5-10 nl of RNase-free water (Invitrogen) was microinjected into one to four animal blastomeres of 4-16-cell embryos. *vangl2* MO (5’-CGTTGGCGGATTTGGGTCCCCCCGA-3’) was described previously ^24^.

RNA or MO-injected embryos were cultured in 0.1x MMR until early gastrula or neurula stages. For a pigmentation assay, embryos were imaged using a Leica stereomicroscope equipped with a color CCD camera. Each experiment included 20-30 embryos per condition. Experiments were repeated at least three times.

### Immunostaining and live time-lapse imaging of Xenopus embryos

For apical domain imaging of neurula, 5 ng of *vangl2* MO-injected or 50 pg of control GFP RNA injected embryos were fixed with MEMFA (100 mM MOPS (pH 7.4), 2 mM EGTA, 1 mM MgSO4, 3.7% formaldehyde)^58^ for 1 hr at room temperature. After permeabilization using 0.1% Triton X-100 in PBS for 10 min, embryos were stained with Alexa Fluor 555-conjugated phalloidin (ThermoFisher Scientific) in PBS containing 1% BSA overnight at 4°C. The dissected neural plate was mounted on a slide glass with two coverglass spacers (0.13-0.17 mm) to minimize damage to the morphology. Embryo images were captured using the LSM980 confocal microscope with a 20x dry objective with 0.55 mm working distance (Zeiss). At least two independent experiments have been done with 20 embryos in each experimental group.

For time-lapse imaging, embryos injected with GFP-Lmo7 RNA or a lineage tracer RNA (myrBFP, myrRFP, GFP-CAAX or 3GFP-ZO1)(100 pg) were cultured until gastrula. Embryos were mounted in 1% low melting temperature agarose (Lonza) on a slideglass attached with a silicone isolator (Grace Bio-labs) or on a glass-bottom dish (Cellvis). Time-lapse imaging was carried out using the AxioZoomV16 fluorescence stereomicroscope (Zeiss) equipped with the AxioCam 506 mono CCD camera (Zeiss) or the LSM880 confocal microscope (Zeiss). Images were taken every 6-10 min for the period of 1.5-4 hrs. Apically constricted (AC) or expanded (AE) cells were defined as the ones, in which the apical domain has been reduced or expanded, respectively, by more than 20%.

### Segmentation, tracking, apical domain assessment

Grayscale images of the cell outline marker were segmented using the Python package Cellpose v2.0.5 ^59^. Each frame was first segmented using the pre-trained model *cyto2* and manually corrected as needed. Cellpose identifies each cell by a pair of non-overlapping connected components: a mask of the cell interior and a mask of the cell outline, with no overlap between different cell outlines. Cell morphology was quantified for each cell interior mask after post-processing steps to ensure tissue confluence. The cell aspect ratio was measured as the ratio between the lengths of the long and short axes of the ellipse with the same normalized second central moments as the cell ^60^. The angle between the long axis and a reference direction defined the cell orientation. Segmented cells were tracked across timepoints using the Python package BayesianTracker (btrack v0.4.5)^61^, where the cell centroid, area, major and minor axes lengths, perimeter, solidity, and higher central moments (up to 3rd order) were used as features in BayesianTracker’s probabilistic model.

For the neural plate segmentation, the neural plate area in stage 14-15 embryos was defined by the brighter phalloidin staining as compared to the surrounding epidermis. Dorsolateral hinge regions were defined as 4-5-cell-wide corridors at the border of the neural plate. The remaining neural plate was defined as “non-hinge” region.

### Statistical analyses

Histograms, dot plots and rose plots for experimental data were generated using the GraphPad Prism 9, PlotsOfData and Rose Diagram Creator, respectively. For cell orientation, the AP axis was set at 0 degrees. The coefficient of variation (CV) and s. d. were caliculated using the GraphPad Prism 9 and used to measure the variability in individual sample groups. The Kolmogorov-Smirnov test was used to determine statistical significance of difference between two groups with non-normal distribution. The Freeman-Halton extension of Fisher’s exact test was used to determine statistical significance of the difference among more than three groups.

### 2D vertex model

Each cell in the 2D vertex model is a polygon. The evolution of the system follows the friction dominated equation of motion for the polygon vertices

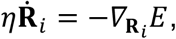

where **R**_*i*_ is the position of vertex *i*, *η* is the viscosity of the vertices, *∇*_**R**_*i*__. indicates the gradient relative to the position of the *i*-th vertex, and *E* is the (dimensionful) energy of the tissue. We use the usual area-and-perimeter-elasticity energy function ^28^:

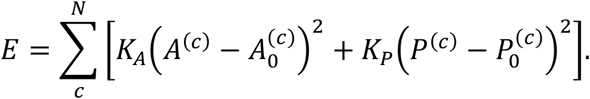

Here, *A*^(*c*)^ and *P*^(*c*)^ are the area and perimeter, respectively, of cell *c*, 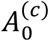 and 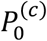 are respectively its target area and perimeter, whereas *K_A_* and *K_P_* are respectively the area and perimeter elasticity moduli. To recast the equation in dimensionless form, we divide this energy by 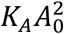, where we assume that the target area of all cells is identical, 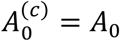. This results in the main text Equation, with the definitions 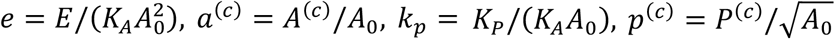, and, 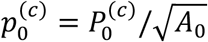. The equation of motions then becomes

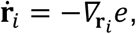

where 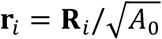 is the dimensionless position of the *i*-th vertex, the overdot indicates the derivative with respect to the dimensionless time (measured in units of *η*/(*K_A_A*_0_)), and the gradient is now with respect to the dimensionless position of vertex *i*.

The initial condition of our model is a regular hexagonal lattice, with *N_x_* + 40 and *N_y_* + 40 cells along the *x* and *y* axis, respectively. Note that every second row contains an extra cell in the neural plate region along the *y* axis, so that the region is symmetrical relative to the AP axis. The outermost cells of the entire model tissue cannot change shape, enforcing fixed boundary conditions. The resulting equations of motion are solved using an explicit Euler scheme, with time step *Δt* = 0.0001, and model tissues are shown at time *t* = 2000. Note that our model does not allow for T1 transitions and the model tissue is therefore always a hexagonal lattice in terms of cell neighbor relations.

### Cell elongation in vertex models

In both 2D and 3D, we define the direction of elongation of a cell as the eigenvector corresponding to the highest eigenvalue of the cell’s gyration tensor

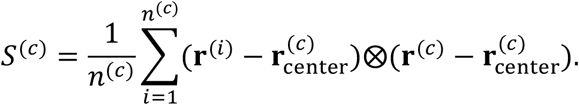

Here, the sum is over the *n*^(*c*)^ vertices corresponding to cell *c*, 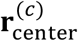 is the average position of all vertices corresponding to that cell, and indicates the outer product. To calculate the elongation of a cell, we use

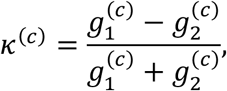

where 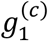 and 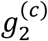 and respectively the first and second largest eigenvalues of *S*^(*c*)^. Lastly, to determine the length and width of the model neural plate region for Fig. 3D, we first select several centrally located cells at each of the outer edges of the model neural plate region. After tissue relaxation, we then measure the distance between the mean centers of relevant pairs of these cell groups.

### 3D vertex model of furrow formation

In the 3D model, cells are represented as polyhedrons with near-constant volume *V_c_* arranged in a mono-layered sheet. The apical and basal sides of the cells are polygons, whereas the lateral sides are quadrilaterals; we use a model based on surface-tensions in cell sides ^62–65^, expanded to also include an apical-perimeter-elasticity term ^66^. The energy function therefore reads

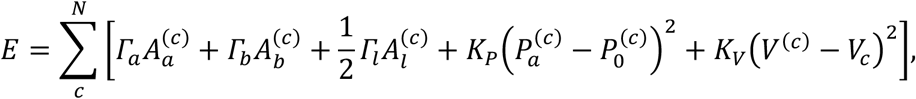

where the sum runs over all *N_c_* cells, 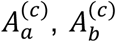, and 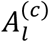 are the apical, basal, and lateral areas of the *i*-th cell, whereas *Γ_a_, Γ_b_* and *Γ_l_* are the corresponding surface tensions and the factor 1/2 arises because each lateral side is shared between two cells. 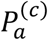 and 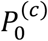 are the current and target apical perimeter of the *i*-th cell, with *K_P_* being the perimeter elasticity modulus. *V*^(*c*)^ is the volume of the *i*-th cell, *V_c_* is the target cell volume, and *K_V_* is the bulk modulus.

To non-dimensionalize the model, we divide it by 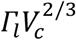, so that the dimensionless energy now reads

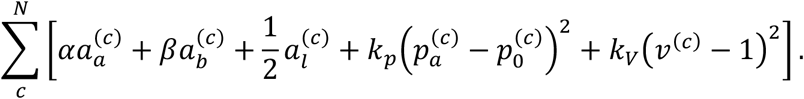

Here, *α* = *Γ_a_*/*Γ_b_* and *β* = *Γ_b_*/*Γ_l_* are the dimensionless apical and basal surface tensions, respectively; 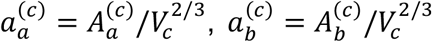, and 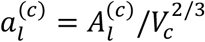 are the apical, basal, and lateral dimensionless areas of the *c*-th cell, respectively; 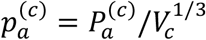 and 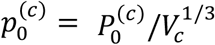 are the dimensionless actual and target apical perimeter of the *c*-th cell, and *v*^(*i*)^ = *V*^(*c*)^/*V_c_* is the dimensionless volume of that cell. The dimensionless perimeter elasticity modulus is given by *k_p_* = *K_P_*/*Γ_l_*, whereas the dimensionless bulk modulus is given by 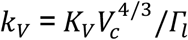. We set *α* = *β* = 0.5, corresponding to a cuboidal tissue, *k_p_* = 1, and *k_V_* = 100. The target apical perimeter is set to equal the equilibrium apical perimeter of a cell without perimeter elasticity in a flat hexagonal honeycomb tissue, 2 · 6^1/3^(*α* + *β*)^−1/3^. To model the constricting cells, we again set their target apical perimeter to equal to 10% of the target perimeter of non-constricting cells. We assume overdamped equations of motion, with dimensionless vertex positions now measured in units of 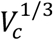 and dimensionless time in units of *η/Γ_l_*. The resulting equations of motion are solved using an explicit Euler scheme, with time step *Δt* = 0.0001; model tissues are shown at *t* = 5000.

The simulations again start with a regular hexagonal lattice. We set boundary conditions along the AP axis to be periodic, as the AP length of the hinge is much larger than its width. Boundary conditions perpendicular to AP are fixed. The simulated patch has 20 cells along the AP axis, and 40 perpendicular to it. As before, the outermost cells perpendicular to AP cannot change shape. Moreover, T1 transitions are again not allowed.

## Acknowledgements

We thank Stas Shvartsman, Rastko Sknepnek, Olga Ossipova, Kyeongmi Kim and the current members of the Sokol laboratory for valuable discussions and Vladimir Gelfand and Mark Corkins for comments on the manuscript. We also wish to thank Matej Krajnc for providing the initial version of the vertex model codes and Damian Dalle Nogare for providing the code for segmentation, tracking and apical domain assessment at the initial stage of this study. We acknowledge the help from the ISMMS Microscopy Core facility. This research was supported by the NIH grants R35GM122492 to SYS and R35GM138380 to KEK. KEK holds an NSF CAREER Award, Packard Fellowship, and Sloan Research Fellowship in Physics. JR acknowledges support from the UK EPSRC (Award EP/W023849/1).

## Supplemental information

**Supplemental Figure 1.**
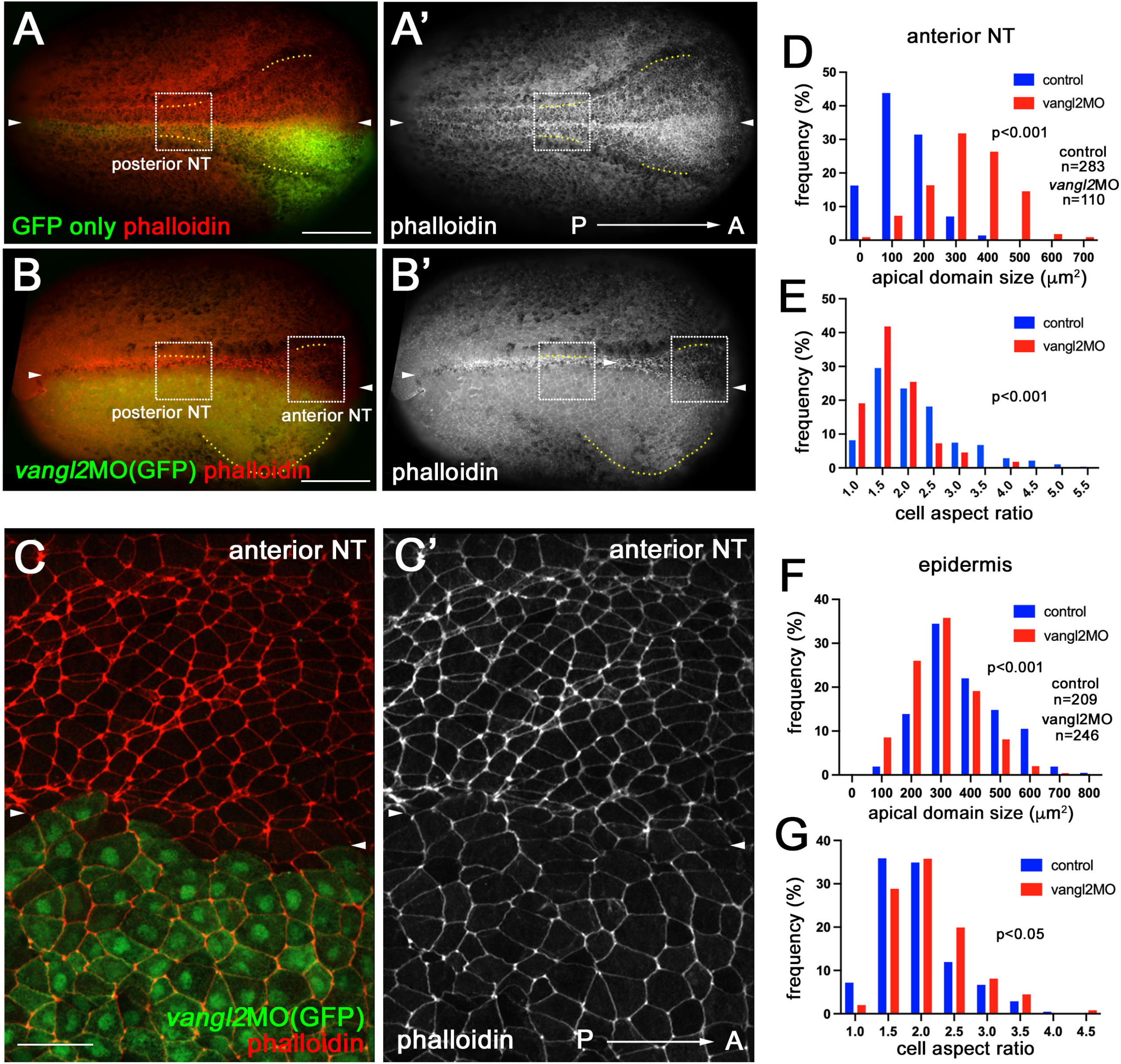
PCP-dependent cell heterogeneity in the anterior and posterior neural plate. (A-G) Vangl2 is required for apical domain heterogeneity (A-B’) Representative images of injected embryos. GFP RNA and *vangl2* MO or control GFP RNA was injected into one dorsal blastomere of 4-cell stage embryos. Embryos at stage 15 were stained with phalloidin. The approximate locations of posterior and anterior NP images in CC’ and Fig. 2A-B’ are indicated in white boxes. The midline is indicated by white arrowheads. (C, C’) *Vangl2* knockdown increased apical domain size and decreased cell aspect ratio in the anterior neural plate. Scale bars are 100 μm in A, B and 50 μm in C. (D) Comparison of apical domain size in *vangl2* MO-injected and control cells, see Methods. (E) Cell aspect ratio for cells in D. (F) Apical domain size in the *vangl2* MO-injected and control epidermis. Note that *vangl2* MO does not enlarge apical domains in the epidermis. (G) Cell aspect ratio in the *vangl2* MO injected and control epidermal cells. N=6 (control). N=9 (*vangl2* MO). Experiments have been repeated three times. Statistical significance was assessed using the Kolmogorov-Smirnov test.

**Supplemental Figure 2.**
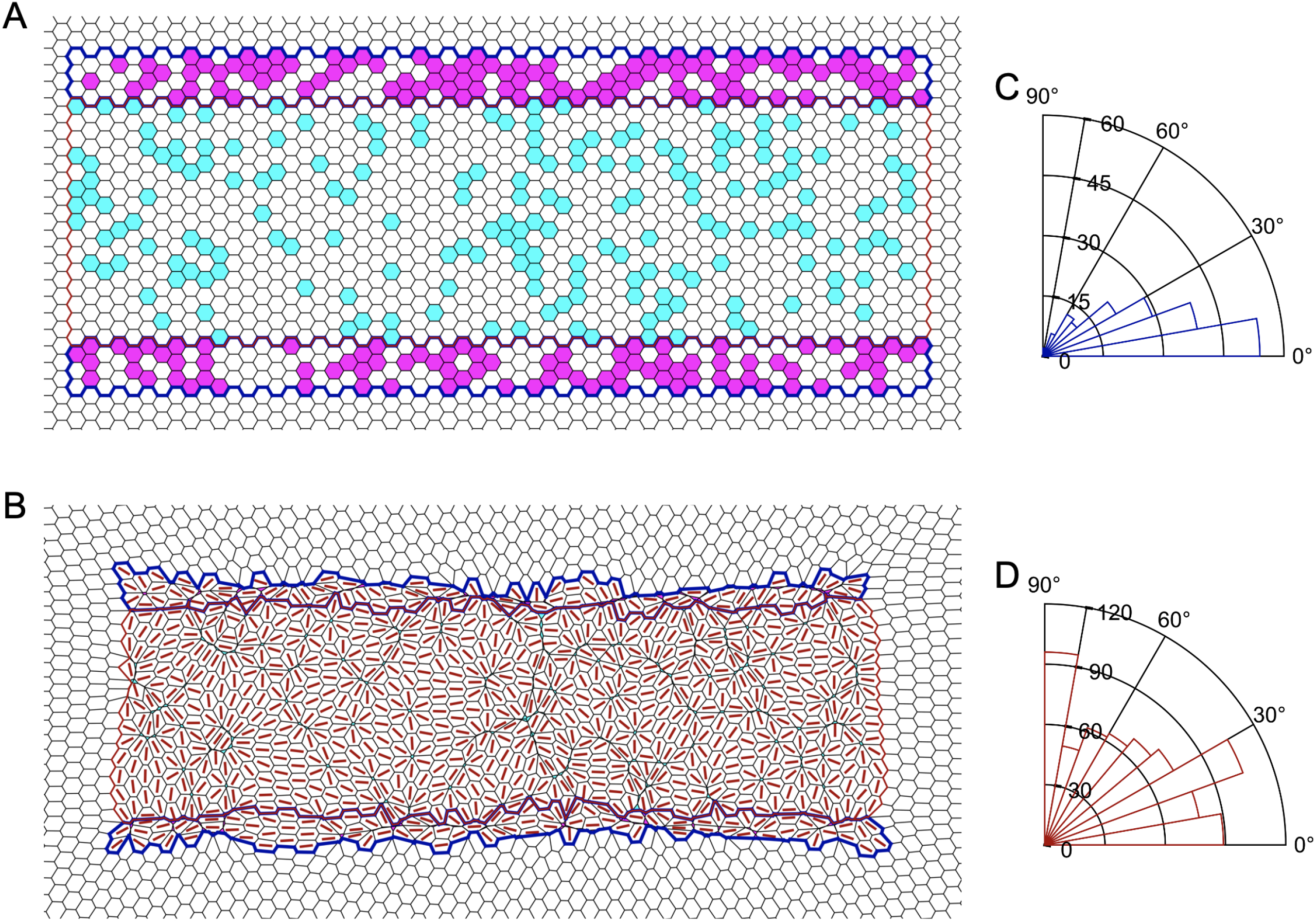
Apically constricting ‘hinge cells’ contribute to neural plate cell morphology and orientation. (A) Zoom on the neural plate region of the vertex model initial condition with separate dorsolateral hinge regions. Blue border outlines the hinges, whereas a red border outlines the region between them. Constricting cells are shown in magenta for the hinges and in cyan for the remainder of the model posterior neural plate (tissue shown for *P_h_* = 0.5 and *P_c_* = 0.2). (B) Model tissue from panel A after relaxation at *t* = 2000. Red lines show the direction of cell elongation. (C-D) Distribution of angles between the AP axis (horizontal line) and cell elongation direction for the model tissue in panel B for non-constricting cells in the hinges (C) and in the remainder of the neural plate (D).

**Supplemental Figure 3.**
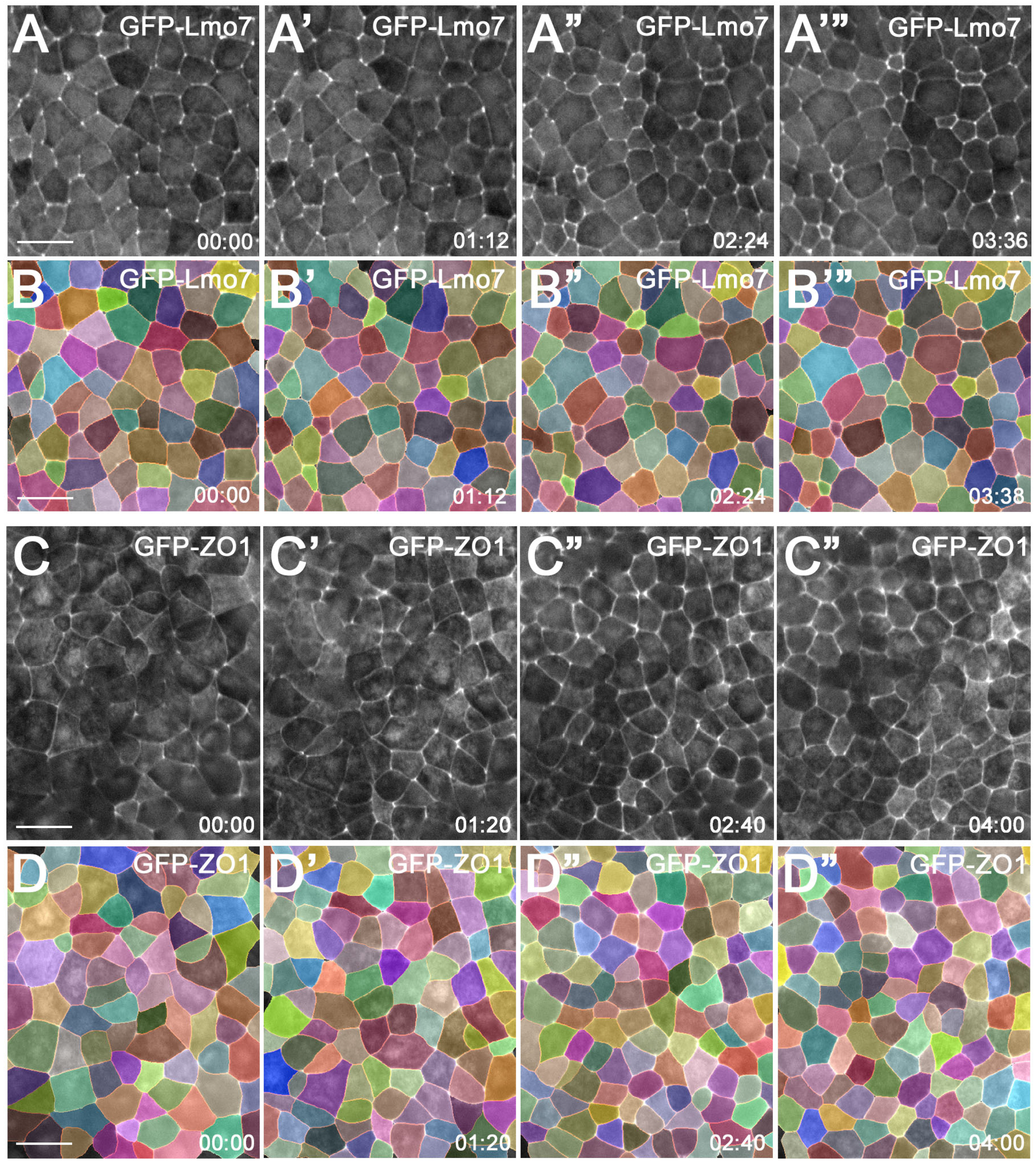
Cell segmentation and cell tracking in Lmo7-expressing and control ectoderm. Representative still images of stage 11-12 ectoderm expressing GFP-Lmo7 (A-A”’) and GFP-ZO1 (C-C”’) from time-lapse imaging. Segmented images of A-A”’ and C-C”’ are shown in B-B”’ and D-D”’, respectively. See Movie 2 and Movie 3. For details of segmentation and cell tracking, see Methods. Scale bars are 40 μm.

**Supplemental Figure 4.**
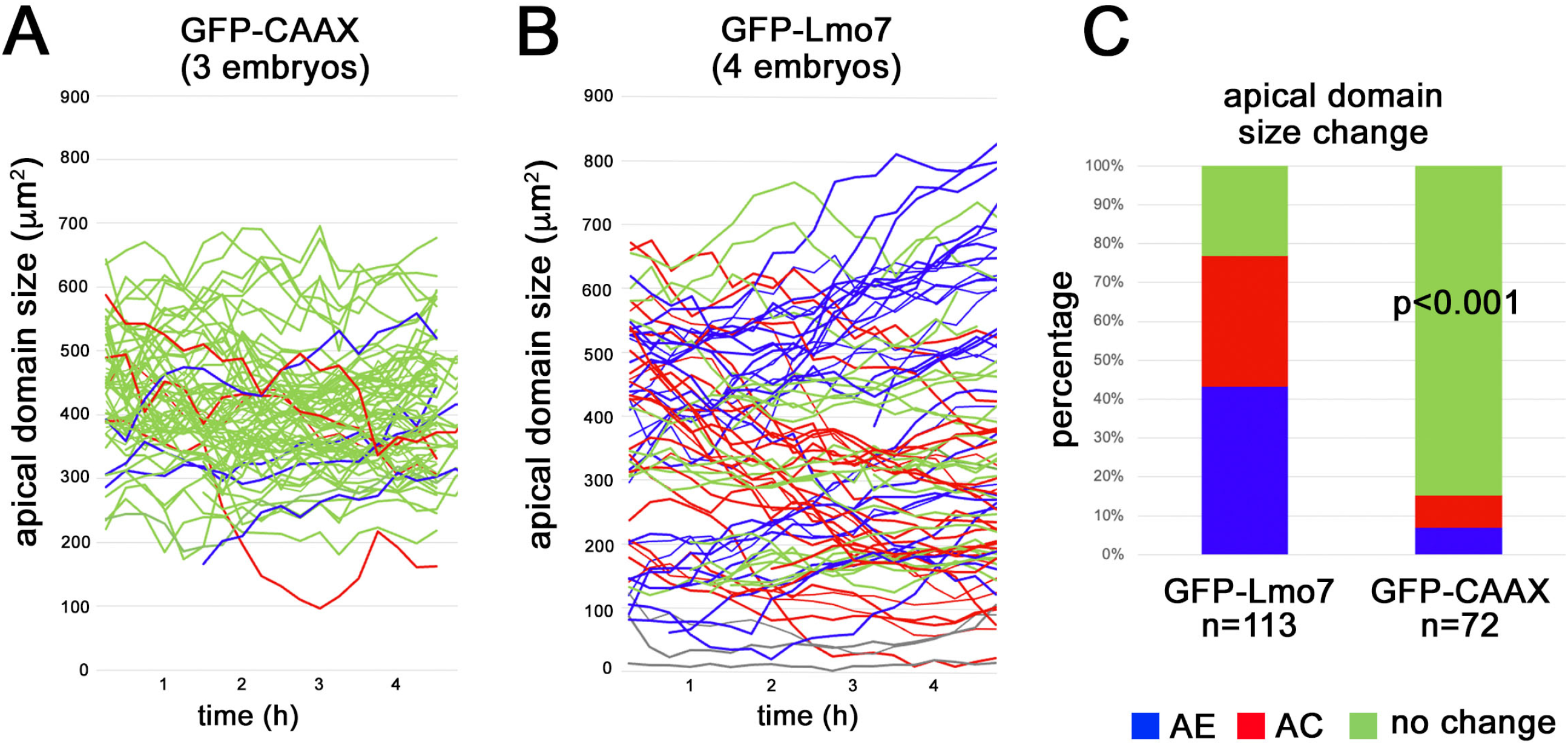
Heterogeneous response of ectodermal cells to Lmo7. (A, B) Quantification of apical domain size dynamics in GFP-CAAX (A) or GFP-Lmo7 (B) embryos. GFP-CAAX or Lmo7 RNA was injected into four animal blastomeres of 4-cell stage embryos as shown in Fig. 5A. Each line represents apical domain size change of an individual cell over 4-5 hours. AC (red) and AE (blue) cells show more than 20% decrease or increase in their apical domain size, respectively. Cells with less than 20% changes (‘no change’) are shown in green. Grey lines indicate cells which had the apical domain smaller than 150 μm^2^ at the beginning of time-lapse imaging and remained small. Data for 72 GFP-CAAX cells are from 3 embryos. Data for 113 GFP-Lmo7 cells are from 4 embryos. (C) Percentage of AC, AE and ‘no change’ cells in A and B. p<0.001. These data have been quantified for 3 embryos in A and 4 embryos in B and represent 3-5 independent experiments. Statistical significance was assessed using the Freeman-Halton extension of Fisher’s exact test.

**Supplemental Figure 5.**
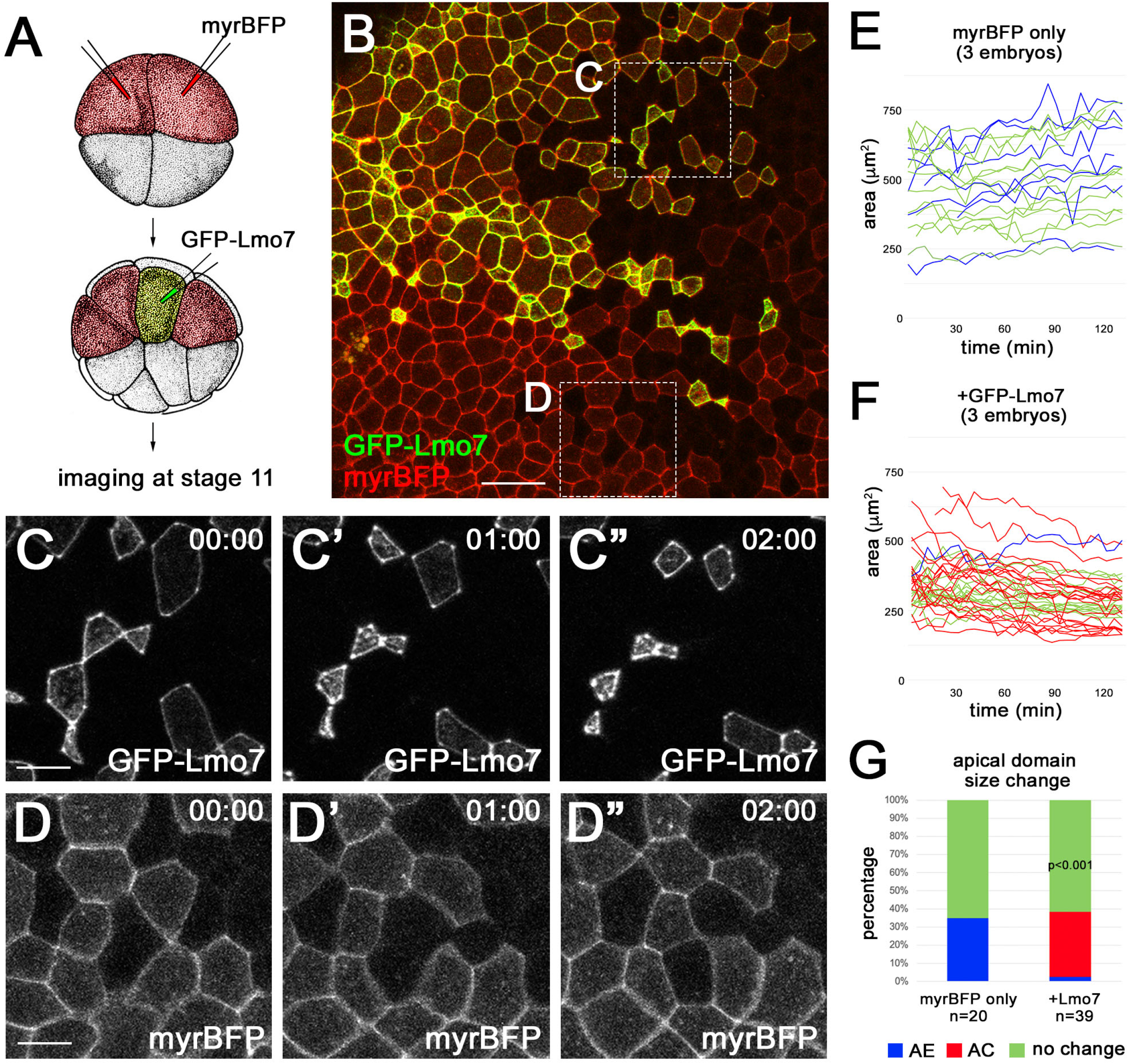
Apical constriction of mosaically expressing Lmo7 cells. (A) Scheme of the experiment. myrBFP RNA (50 pg) was injected into two ventral blastomeres of 4-cell stage embryos. GFP-Lmo7 RNA (200 pg) was subsequently injected into one ventral blastomere of 16-cell stage embryos. The animal ectoderm was imaged at stage 11. (B) Representative image of myr-BFP and GFP-Lmo7 expressing embryos. (C-D”) Representative still images of GFP-Lmo7 (C-C”) and myrBFP (D-D”) cells from time-lapse imaging. (E, F) Quantification of apical domain size dynamics in myrBFP (E) and GFP-Lmo7 (F) cells. Each line represents apical domain size changes of individual cells over time. n=20 cells (myrBFP only) and n=39 (+GFP-Lmo7) from three embryos. (G) Percentage of AC (red), AE (blue) and ‘no change’ (green) cells (See Fig. 5 legend) is shown in E and F. Statistical significance, p<0.001; the Freeman-Halton extension of Fisher’s exact test. The data represent 3-5 independent experiments.

**Supplemental Figure 6.**
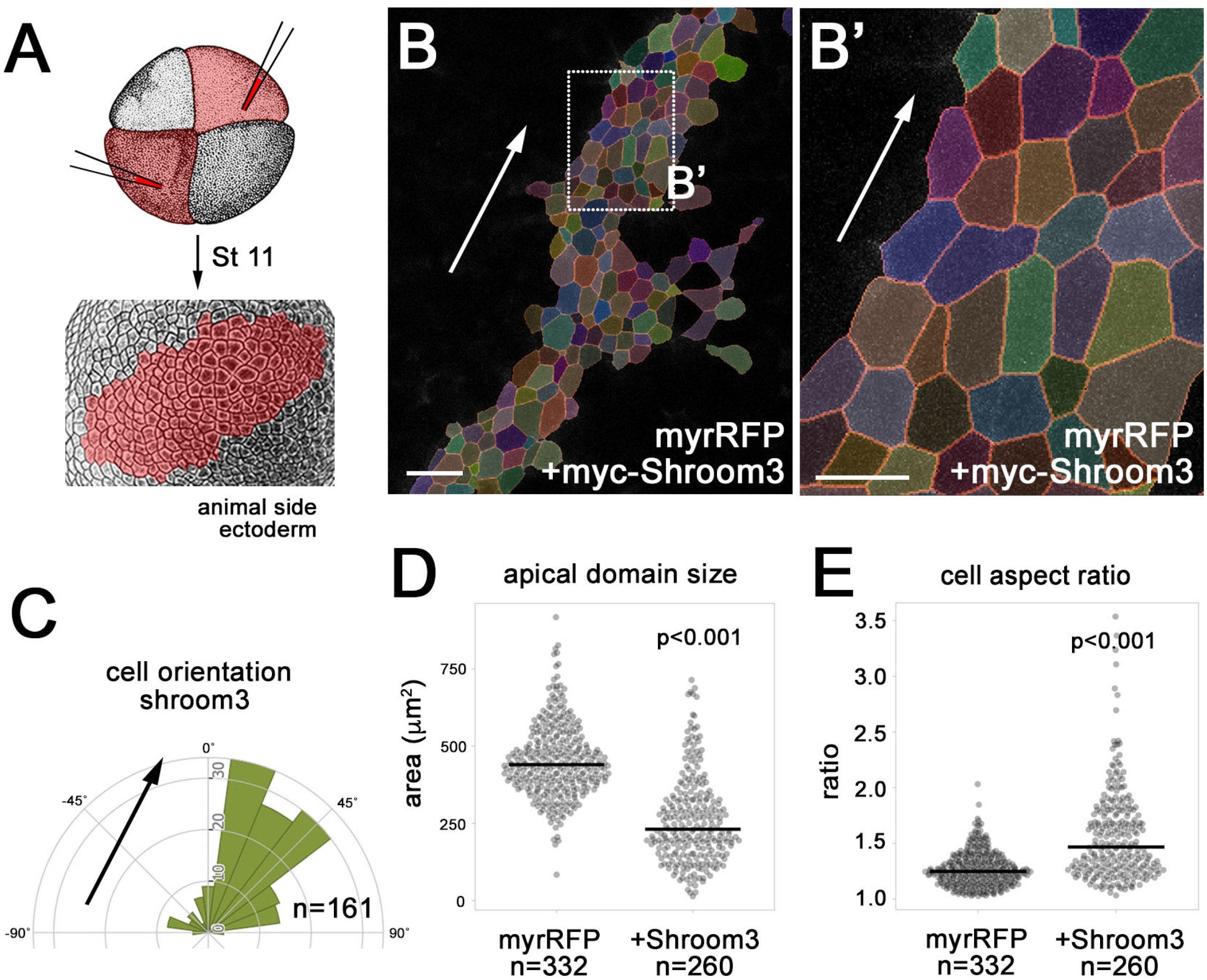
Shroom3 promotes apical domain heterogeneity. (A) Scheme of the experiment for B-E. myc-Shroom3 (40 pg) and myrRFP (50 pg) RNAs were co-injected into two opposing animal blastomeres of 4-8 cell stage embryos. Animal pole ectoderm was imaged at stage 11. (B, B’) Representative segmented Shroom3-expressing ectoderm. Rectangular area in B is enlarged in B’. (C) Rose plot depicting cell orientation of Shroom3 cells in B and B’. n=161. (D, E) Quantification of apical domain size (D) and cell aspect ratio (E) of myrRFP cells and myrRFP+Shroom3 cells. n=332 (myrRFP) and n=260 (+Shroom3). Two embryos each. p<0.001, Kolmogorov-Smirnov test. Scale bars are 50 μm in B and 20 μm in B’.

**Supplemental Table 1.** Coefficients of variation for data shown in Fig. 1 and Fig. 2.

**Coefficient of variation (CV)**
|  | **NT hinge** | **NT non-hinge** | **st 11 ectoderm** |
| --- | --- | --- | --- |
| apical domain size | 60.59% | 42.80% | 31.79% |
| cell aspect ratio | 50.50% | 34.55% | 20.51% |

|  | **control** | ***vangl2* MO** |
| --- | --- | --- |
| apical domain size | 73.07% | 37.99% |
| cell aspect ratio | 42.40% | 25.21% |

**Supplemental Movie 1. Ectoderm morphology in embryos expressing GFP-Lmo7.**

Time-lapse recording of GFP-Lmo7 expressing cells in stage 11 *Xenopus* ectoderm. 100 pg of GFP-Lmo7 RNA was injected into the two animal blastomeres of 4-cell stage embryos. Animal side view. Images were taken every 10 min on the Zeiss AxioZoom stereo microscope. Duration is 3 hrs and 20 min.

**Supplemental Movie 2. Apical domain dynamics of GFP-Lmo7 cells.**

Time-lapse recording of GFP-Lmo7 expressing cells in stage 11 *Xenopus* ectoderm. 100 pg of GFP-Lmo7 RNA was injected into four animal blastomeres of 4-cell stage embryos to obtain uniform expression. Animal view. Images were taken every 10 min on the Zeiss AxioZoom stereo microscope. Duration of the time-lapse video is 4 hrs.

**Supplemental Movie 3. Apical domain dynamics of embryonic ectoderm cells expressing ZO1.**

Time-lapse recording of control 3xGFP-ZO1 expressing cells in stage 11 *Xenopus* ectoderm. 100 pg of 3xGFP-ZO1 RNA was injected into four animal blastomeres of 4-cell stage embryos to obtain uniform expression. Animal view. Images were taken every 8 min on the Zeiss AxioZoom stereo microscope. Duration of the time-lapse video is 3 hrs and 36 min.

**Supplemental Movie 4. Apical domain dynamics of embryonic ectoderm cells uniformly expressing Lmo7.**

Time-lapse recording of cells co-expressing 3xGFP-ZO1 and Flag-Lmo7 in stage 11 *Xenopus* ectoderm. 100 pg of 3xGFP-ZO1 and 100 pg of Flag-Lmo7 RNA was injected into four animal blastomeres of 4-cell stage embryos to obtain uniform “sheet” expression. Animal view. Images were taken every 6 min on the Zeiss LSM880 confocal microscope. Duration is 1 hr and 42 min.

**Supplemental Movie 5. Apical domain dynamics of ectoderm in embryos mosaically expressing Lmo7.**

Time-lapse recording of cells co-expressing 3xGFP-ZO1 and Flag-Lmo7 in stage 11 *Xenopus* ectoderm. 100 pg of 3xGFP-ZO1 and 100 pg of Flag-Lmo7 RNA were injected into one animal blastomere of 4-cell stage embryos to obtain “mosaic” expression. Animal view. Images were taken every 6 min on the Zeiss LSM880 confocal microscope system. Duration is 1 hr 54 min.

